# Low-background Acyl-biotinyl Exchange Largely Eliminates the Co-isolation of Non-S-acylated Proteins and Enables Deep S-acylproteomic Analysis

**DOI:** 10.1101/588988

**Authors:** Bo Zhou, Yang Wang, Yiwu Yan, Javier Mariscal, Dolores Di Vizio, Michael R. Freeman, Wei Yang

## Abstract

Protein *S*-acylation (also called palmitoylation) is a common post-translational modification whose deregulation plays a key role in the pathogenesis of many diseases. Acyl-biotinyl exchange (ABE), a widely used method for the enrichment of *S*-acylated proteins, has the potential of capturing the entire *S*-acylproteome in any types of biological samples. Here, we showed that current ABE methods suffer from high background arising from the co-isolation of non-*S*-acylated proteins. The background can be substantially reduced by an additional blockage of residual free cysteine residues with 2,2’-dithiodipyridine prior to biotin-HPDP reaction. Coupling the low-background ABE (LB-ABE) method with label-free quantitative proteomics, 2,895 high-confidence candidate *S*-acylated proteins (including 1,591 known *S*-acylated proteins) were identified from human prostate cancer LNCaP cells, representing so-far the largest *S*-acylproteome dataset identified in a single study. Immunoblotting analysis confirmed the *S*-acylation of five known and five novel prostate cancer-related *S*-acylated proteins in LNCaP cells and suggested that their *S*-acylation levels were about 0.6-1.8%. In summary, the LB-ABE method largely eliminates the co-isolation of non-*S*-acylated proteins and enables deep *S*-acylproteomic analysis. It is expected to facilitate much more comprehensive and accurate quantification of *S*-acylproteomes than previous ABE methods.

---

Protein *S*-acylation (more commonly known as protein *S*-palmitoylation or simply palmitoylation) is a protein post-translational modification, where long-chain fatty acids (predominantly the 16-carbon palmitic acid) are covalently attached to cysteine residues via thioester bonds.^1^ Unlike other lipid modifications such as myristoylation and prenylation, protein *S*-acylation is reversible. The cycling between the *S*-acylation and de-*S*-acylation states dynamically regulates protein localization, trafficking, activity, stability, and complex formation.^2^ Therefore, protein *S*-acylation plays a key role in various biological processes such as signal transduction and metabolism, and its aberration leads to many human diseases such as cancer and neurodegenerative disorders.^3^

Traditionally, protein *S*-acylation is almost exclusively analyzed using [^3^H]-palmitate metabolic labeling followed by immunoprecipitation and days to weeks of film exposure, on the basis of individual proteins.^4^ To enable the analysis of protein *S*-acylation at the proteome level, two complementary approaches (*i.e.*, palmitate-centric and cysteine-centric) were developed.^5–7^ In the palmitate-centric approach, cells are metabolically labeled with an azido- or alkyne-modified fatty acid probe. The probe-modified proteins are then conjugated to biotin by click chemistry, so they can be enriched by streptavidin affinity purification and analyzed by liquid chromatography-tandem mass spectrometry (LC-MS/MS).^8^ Thus, the palmitate-centric approach was also termed <u>m</u>etabolic labeling with a palmitic acid analog followed by <u>c</u>lick <u>c</u>hemistry (MLCC).^9^ Although particularly useful for investigating *S*-acylation dynamics in cultured cells,^10^ MLCC suffers from several inherent limitations: a) it cannot typically be used to analyze tissue and biofluid samples because of a dependence on metabolic labeling, b) it needs optimization for different cell types to achieve best sensitivity and to minimize cell death, c) it is biased towards the enrichment of *S*-acylated proteins with rapid turnover, d) it provides a relatively low recovery of *S*-acylated proteins, especially in cells with high lipogenic activity, where *S*-acylated proteins are mostly labeled by native palmitate, and e) the intact probe-modified peptides are difficult to analyze by LC-MS/MS due to their high hydrophobicity.^5^

In complement to MLCC, a metabolic labeling-independent, cysteine-centric approach named acyl-biotinyl exchange (ABE) was also developed for the enrichment of *S*-acylated proteins.^11^ In ABE, free cysteine residues are blocked by an alkylation reagent, *S*-acyl groups are specifically removed from cysteine residues by using neutral hydroxylamine (Hyd) to cleave thioester bonds, and the newly formed thiol groups are labeled with biotin-HPDP. As such, *S*-acylated proteins are converted into biotinylated proteins, so they can be enriched by streptavidin affinity purification and identified by LC-MS/MS.^12^ Different from the MLCC, the ABE approach can be used to analyze *S*-acylated proteins in all types of biological samples and has the potential of capturing the entire *S*-acylproteome.^5^

Previously, we reported an ABE-derived method termed Palmitoyl-protein Identification and Site Characterization (PalmPISC)^13^. In PalmPISC, disulfide bonds are reduced by tris(2-carboxyethyl)phosphine (TCEP), and non-*S*-acylated cysteine residues are irreversibly blocked by N-ethylmaleimide (NEM). Subsequently, *S*-acylated cysteine residues are converted by neutral Hyd into free cysteine residues, which are further biotinylated by biotin-HPDP. As a control, Tris buffer is used to replace neutral Hyd. The *in vitro* biotinylated (formerly *S*-acylated) proteins are enriched by streptavidin affinity purification, selectively eluted by TCEP, and analyzed by LC-MS/MS. Proteins significantly enriched in the Hyd(+) condition, compared with the control Hyd(-) condition, are considered as candidate *S*-acylated proteins. Using PalmPISC, we identified 398 *S*-acylated proteins (out of 928 identified proteins) from human prostate cancer (PCa) DU145 cells,^13^ as well as 215 *S*-acylated proteins (out of >1,300 identified proteins) from human platelets.^14^

Owing to their advantages, ABE methods have been widely used in protein *S*-acylation and *S*-acylproteomic studies. Nevertheless, we and others found that a significant amount of proteins was repeatedly obtained in the control (*i.e.*, Hyd−) condition, accounting for about 30-80% amount of proteins enriched in the experimental (*i.e.*, Hyd+) condition. The background (*i.e.*, co-isolated non-*S*-acylated proteins) causes at least two major concerns. Firstly, some co-isolated non-*S*-acylated proteins can be several orders of magnitude more abundant than low-abundance *S*-acylated proteins, and thus mask their detection and identification by LC-MS/MS. Secondly, for a specific *S*-acylated protein, a fraction of its non-*S*-acylated form might be co-isolated with the *S*-acylated form, resulting in an underestimation of actual quantitative ratios between different biological conditions, a phenomenon known as “ratio compression”. In quantitative *S*-acylproteomic studies, different *S*-acylproteins may suffer from different levels of ratio compression; hence, their quantification accuracy is compromised to different degrees.

The present study aimed to address the following pressing questions: 1) To what extent do ABE methods suffer from the background limitation? 2) Can we substantially reduce the background and improve the signal-to-noise ratios for enriched *S*-acylated proteins? 3) Will a low-background ABE (LB-ABE) method enable deep *S*-acylproteomic analysis by discovering more low-abundance *S*-acylated proteins? Here, we showed that 1) only ∼35% of identified proteins were enriched by ≥ 1.5-fold (Hyd+/Hyd−) by regular ABE and could be considered as candidate *S*-acylated proteins, 2) using 2,2’dithiodipyridine (DTDP) to further block residual free cysteine residues that are not alkylated by NEM can substantially decrease the background, and 3) the LB-ABE method enabled the identification of ∼1,600 known and ∼1,300 novel candidate human *S*-acylated proteins from human PCa LNCaP cells, of which 10 proteins were validated by immunoblotting analysis. The LB-ABE method and the comprehensive *S*-acylproteomic dataset are expected to be valuable for the fields of *S*-acylation, proteomics, and cancer research.

## EXPERIMENTAL SECTION

### Materials

Iodoacetamide was purchased from GE Healthcare. RPMI-1640 and DMEM media were from Gibco. Microcon YM-30 spin filters were from Millipore. MS-grade Trypsin, Asp-N, and Arg-C were from Promega. Tris, SDS, methanol, chloroform, 2,2’-dithiodipyridine (DTDP), dimethylformamide (DMF), hydroxylamine, sodium chloride, ammonium bicarbonate, and ammonium formate were from Sigma-Aldrich. Arginine (Arg0), lysine (Lys0), ^13^C_6_^15^N_4_-arginine (Arg10), ^13^C_6_^15^N_2_-lysine (Lys8), dialyzed fetal bovine serum, RPMI 1640 medium for SILAC, 660 nm Protein Assay Kit, tris(2-carboxyethyl)-phosphine (TCEP), N-ethylmaleimide (NEM), high-capacity streptavidin agarose beads, SuperSignal chemiluminescent substrate, trap columns, EASY-Spray analytical columns, and Horseradish peroxidase (HRP)-conjugated streptavidin were from Thermo Fisher Scientific. Primary antibodies were from Cell Signaling Technology, Novus Biologicals, Santa Cruz Biotechnology, Sigma-Aldrich, and R&D systems. HRP-conjugated species-specific secondary antibodies were from Cell Signaling Technology and Jackson ImmunoResearch Laboratory.

### Cell Culture

Human prostate cancer LNCaP, DU145, and PC3 cells were obtained from the American Type Culture Collection and authenticated by the Laragen Inc. using the Promega PowerPlex 16 system. Cells were free of mycoplasma contamination as determined using the MycoAlert PLUS Mycoplasma Detection Kit (Lonza) by following the manufacturer’s protocol. For SILAC labeling, LNCaP cells were grown for three passages (∼8 doublings) in Arg/Lys-depleted RPMI-1640 medium supplemented with 10% (v/v) dialyzed fetal bovine serum and Arg0/Lys0 or Arg10/Lys8, essentially as we previously described.^15,16^ When cells grew to 80-90% confluency, they were harvested with cell scrapers, washed with cold phosphate buffered saline for three times, and pelleted by centrifugation at 500 × *g* for 5 min at 4°C.

### ABE and LB-ABE Enrichment

The ABE enrichment was performed essentially as we previously described in the PalmPISC method with minor modifications.^13^ Briefly, after cell lysis, proteins were reduced with 50 mM TCEP, alkylated with 50 mM NEM, biotinylated with 1 mM biotin-HPDP in the presence or absence of 2 M Hyd, enriched by streptavidin affinity purification, eluted by 50 mM TCEP, and precipitated by methanol/chloroform precipitation. For LB-ABE enrichment, reaction with 25 mM DTDP was performed between NEM alkylation and biotin-HPDP reaction.

### UHPLC-MS/MS Analysis

Proteins were digested with trypsin, Asp-N, or Arg-C by filter-aided sample preparation (FASP)^17^ or with trypsin in gel as we previously described.^18^ The resulting peptides were separated by EASY-Spray columns and analyzed by an LTQ Velos, an LTQ Orbitrap Elite, or an Orbitrap Fusion Lumos mass spectrometer essentially as described.^19,20^ Database searching analyses were conducted using MaxQuant^21^ and Andromeda^22^. Data analysis was performed using Perseus^23^ and R (www.R-project.org). The mass spectrometry proteomics data have been deposited to the ProteomeXchange Consortium (http://proteomexchange.org) via the PRIDE partner repository^24^ with database identifiers PXD013187 and PXD013189.

### Immunoblotting Analysis

Immunoblotting analysis was performed as described.^13,25^ Briefly, proteins were separated by SDS-PAGE and electro-transferred onto PVDF membranes. Membranes were blocked with TBS, 0.1% Tween 20, 5% non-fat milk for 1 h at RT, incubated with primary antibodies overnight at 4°C, followed by incubation with species-specific HRP-conjugated secondary antibodies for 1 h at RT. Signals were detected using SuperSignal chemiluminescent reagent followed by exposure of blots to X-ray films. Densitometric analysis was performed using ImageJ (v1.52a).

More detailed experimental procedures can be found in the Supporting Information.

## RESULTS AND DISCUSSION

### Current ABE Methods Suffer from High Background

So far, many global *S*-acylproteomic studies were performed using ABE and its derived methods, which are independent of metabolic labeling and have the potential of capturing the entire *S*-acylproteome in any biological samples including tissues and extracellular vesicles in biofluids.^5,26^ In theory, ABE methods are able to capture more *S*-acylated proteins than other methods such as MLCC, which is biased towards the enrichment of *S*-acylated proteins with rapid turnover.^5^ Nevertheless, an analysis of *S*-acylated proteins compiled in the SwissPalm (v2)^3^―a comprehensive database of *S*-acylated proteins identified since 1981―revealed that only 41% (1,173 out of 2,881) of the human *S*-acylproteome and 33% (976 out of 2,939) of the mouse *S*-acylproteome were discovered by ABE studies (Fig. 1; Tables S1 and S2). This indicates that the current ABE methods suffer from certain limitations and need substantial improvement to realize their full potential of deep and unbiased *S*-acylproteomic analysis.

**Figure 1.**
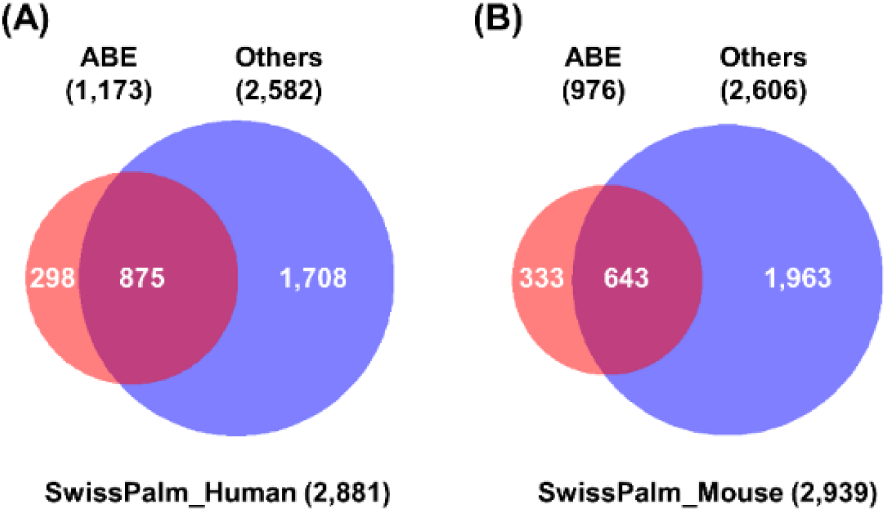
Venn diagrams of **(A)** human and **(B)** mouse *S*-acylated proteins identified by ABE and other methods. The numbers were retrieved from the SwissPalm (v2) database.

Previously, our review of published *S*-acylproteomic studies identified a common problem with ABE methods: significant contamination with non-*S*-acylated proteins, which hinders the detection of lowly abundant *S*-acylated proteins and causes ratio compression in quantitative *S*-acylproteomic comparison studies.^5^ Nonetheless, the extent of such background noise (*i.e.*, co-isolated non-*S*-acylated proteins) remains not well defined because most ABE-based *S*-acylproteomics studies applied spectral counting,^27^ a semi-quantitative proteomic approach, to distinguish candidate *S*-acylated proteins from co-isolated non-*S*-acylated proteins.^5^

To quantitatively assess the background level, stable isotope labeling by amino acid in cell culture (SILAC),^28^ a highly accurate quantitative proteomics method, was coupled with our PalmPISC method to analyze the *S*-acylproteome of human prostate cancer (PCa) LNCaP cells (Fig. 2A). Established in 1980, LNCaP has been one of the most widely used cell lines in PCa research.^29,30^ Proteins extracted from light SILAC-labeled cells were treated with neutral hydroxylamine (Hyd) to de-*S*-acylate proteins, whereas proteins extracted from heavy SILAC-labeled cells were treated with control Tris-HCl buffer. Subsequently, light and heavy proteins were mixed and subjected to ABE enrichment using our PalmPISC approach. In-gel tryptic digestion followed by liquid chromatography-tandem mass spectrometry (GeLC-MS/MS) analysis identified a total of 1,654 proteins with a false discovery rate (FDR) of ≤1%, of which 1,571 proteins were quantified (Table S3). Using the empirical cutoff of heavy/light ratio of 1.5, 552 (35.1%) and 1,019 (64.9%) proteins were respectively considered as candidate *S*-acylated proteins (ratio≥1.5) and non-*S*-acylated proteins (ratio<1.5) (Table S3). Figure 2B shows representative SILAC spectra of candidate *S*-acylated proteins (upper panel) and co-isolated contaminant proteins (lower panel). A density plot analysis of the 1,571 quantified proteins suggested that most proteins have SILAC ratios of <1.5, of which many have SILAC ratios of about 1 (*i.e.*, log_2_-ratios of about 0) (Fig. 2C). In addition, many co-isolated non-*S*-acylated proteins have total ion intensities comparable to those of *S*-acylated protein candidates (Fig. 2D). The median log_2_-transformed total ion intensity of contaminant proteins is only slightly lower than that of candidate *S*-acylated proteins (19.6 *vs* 20.7) (Fig. 2E). Moreover, consistent with our finding, two other *S*-acylproteomic studies using SILAC and different ABE-derived methods also showed that about 65% of identified proteins were non-*S*-acylated proteins (Fig. 2F).^9,31^ Collectively, the quantitative analyses suggested that the current ABE methods generally suffer from high background.

**Figure 2.**
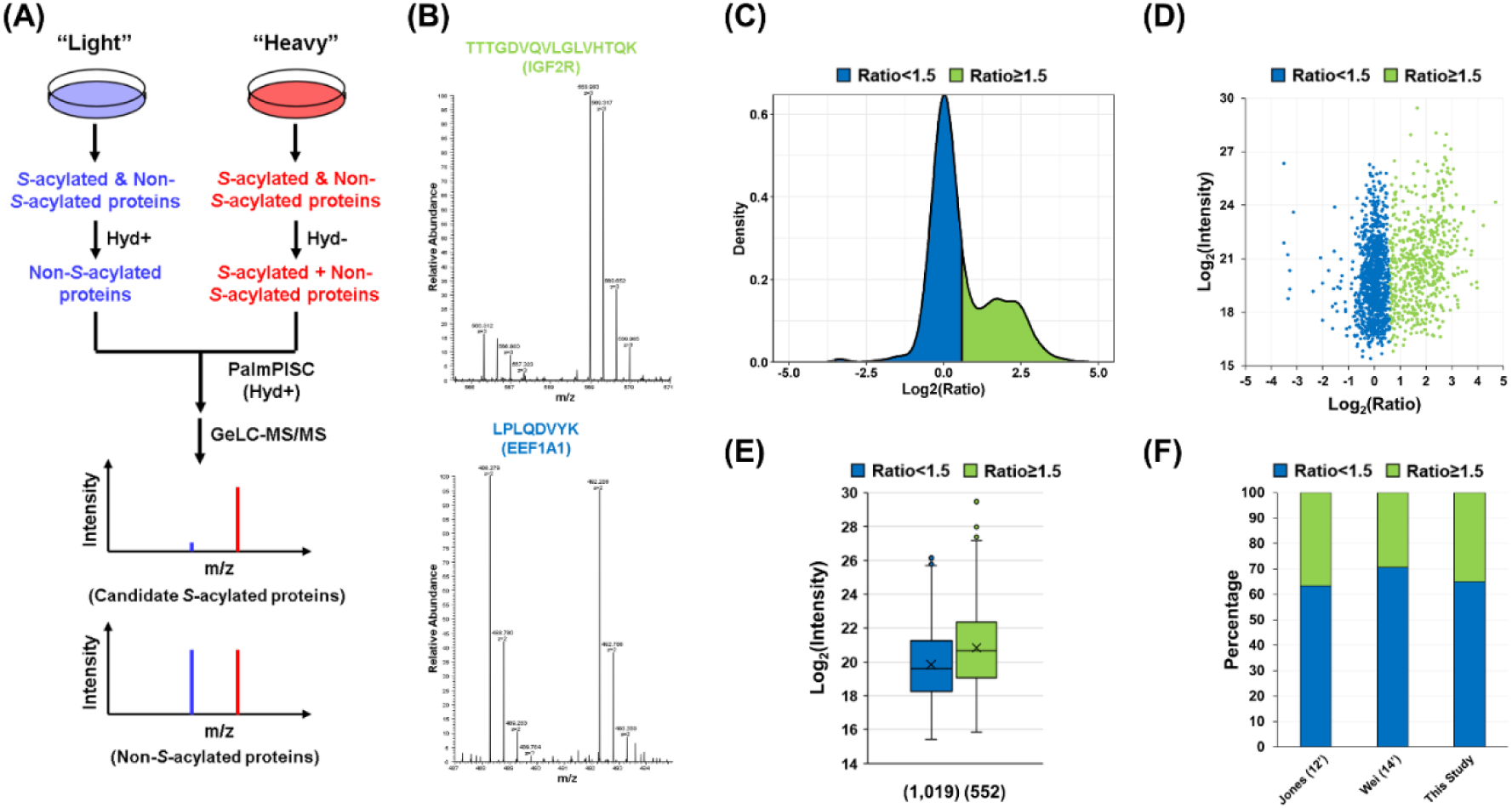
SILAC analysis to determine the extent of contamination caused by the co-isolation of non-*S*-acylated proteins. **(A)** Workflow of SILAC analysis to distinguish candidate *S*-acylated proteins from non-*S*-acylated proteins. Hyd: hydroxylamine. **(B)** Representative SILAC pairs showing the identification of candidate *S*-acylated proteins (upper) and non-*S*-acylated proteins (lower). **(C)** Density plot of log_2_-transformed heavy/light SILAC ratios. The ratio of 1.5 corresponds to log_2_ratio of 0.585. **(D)** Scatter plot of log_2_-transformed heavy/light SILAC ratios against log_2_-transformed total ion intensities. **(E)** Box-plot comparison of log_2_-transformed total ion intensities of non-*S*-acylated proteins (SILAC ratio<1.5) and candidate *S*-acylated proteins (SILAC ratio≥1.5). **(F)** Stacked histogram showing the percentages of non-*S*-acylated proteins in three independent SILAC-and ABE-based *S*-acylproteomic profiling studies.^9,31^

### Blockage of Residual Free Cysteine Residues with 2,2’-dithiodipyridine (DTDP) Largely Eliminates the Co-isolation of Non-*S*-acylated Proteins

The presence of a high background of contaminant proteins may severely mask the detection of low-abundance *S*-acylated proteins and compress the signal-to-noise ratios of many *S*-acylated proteins, especially those of low *S*-acylation levels. Consequently, this not only hinders deep *S*-acylproteomic profiling but also decreases the accuracy of quantifying *S*-acylated proteins across different samples. Thus, it is important to eliminate the co-isolation of non-*S*-acylated proteins. After much trial and error, we reasoned that the co-isolation of non-*S*-acylated proteins is likely caused by the reaction of biotin-HPDP with certain residual free cysteine residues that, due to some unknown reasons, cannot be completely blocked by NEM even with longer incubation time. Therefore, after NEM alkylation and prior to biotin-HPDP reaction, use of DTDP (a thiol-reactive reagent similar to HPDP) to further block these residual cysteine residues may prevent their biotinylation by biotin-HPDP and thus largely eliminate the co-isolation of non-*S*-acylated proteins (Fig. 3).

**Figure 3.**
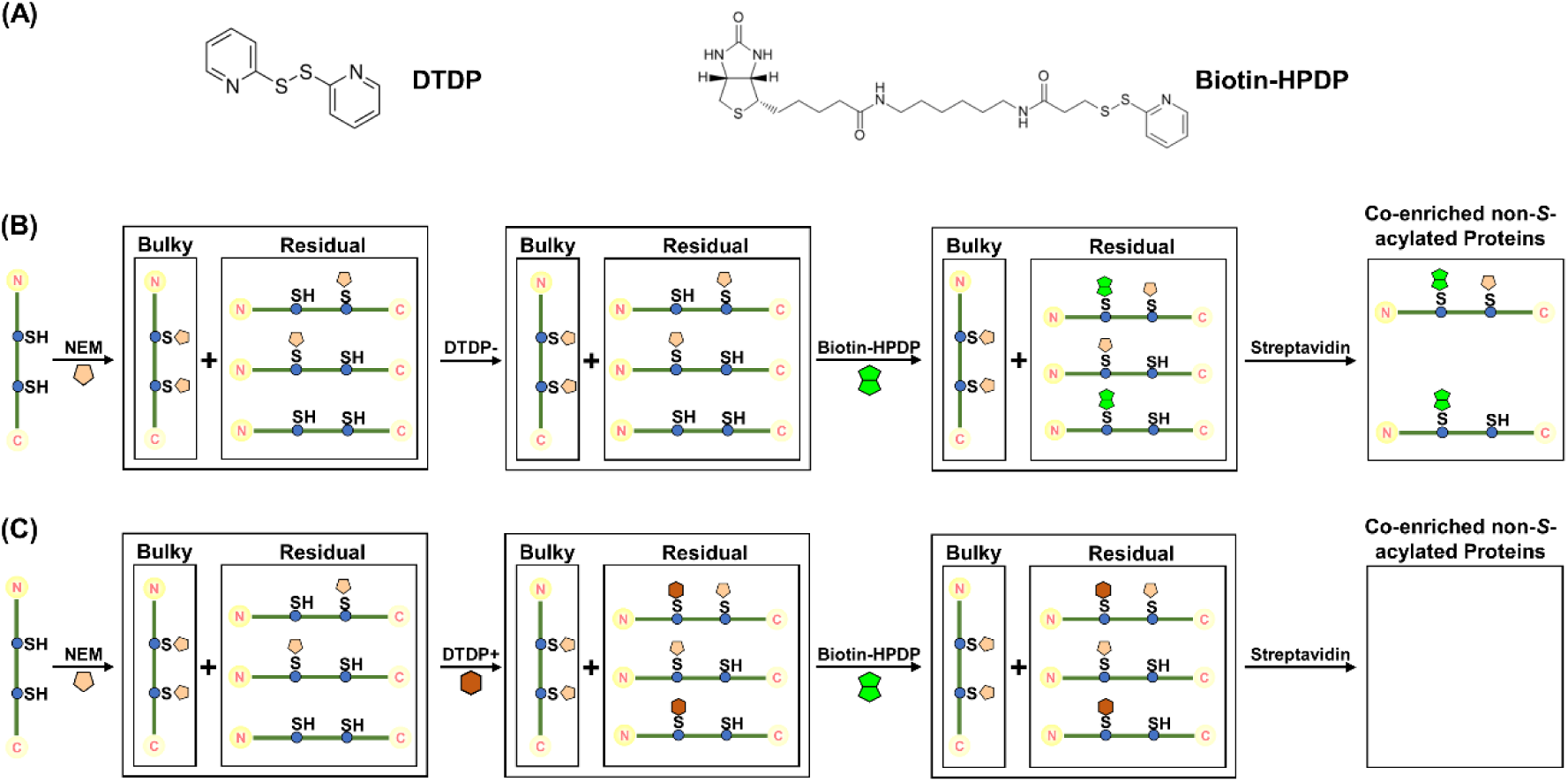
Potential mechanisms for the co-enrichment of non-*S*-acylated proteins by ABE and its prevention by DTDP reaction. **(A)** Structural comparison of DTDP and biotin-HPDP. **(B)** Potential mechanism for the co-enrichment of non-*S*-acylated proteins. After NEM alkylation, certain (but not all) residual free cysteine residues may react with biotin-HPDP, resulting in the co-isolation of proteins containing such cysteine residues. **(C)** Potential mechanism for the elimination of co-enrichment of non-*S*-acylated proteins by DTDP reaction. DTDP may block the residual HPDP-reactive free cysteine residues and thus prevents their biotinylation by biotin-HPDP. Consequently, non-*S*-acylated proteins containing these cysteine residues will not be co-enriched by streptavidin affinity purification. Green lines: cysteine-containing proteins; SH: thiol groups of cysteine residues; S: cysteine sulfur.

Indeed, with an additional alkylation step using DTDP (Fig. 4A), the level of co-enriched non-*S*-acylated proteins from human PCa DU145 whole cell lysates was significantly (*p*=0.003) reduced from 33.8% (± 2.5%) to 9.8% (±0.4%) (*i.e.*, 3.4-fold reduction) (Fig. 4B). Hereinafter, we termed the new method containing the DTDP reaction step “low-background ABE” (LB-ABE). Using the LB-ABE method, lowbackground enrichment of *S*-acylated proteins was obtained by a different operator using human PCa PC3 cells (Fig. S1), confirming that the method is cell line/operator-independent and reproducible.

**Figure 4.**
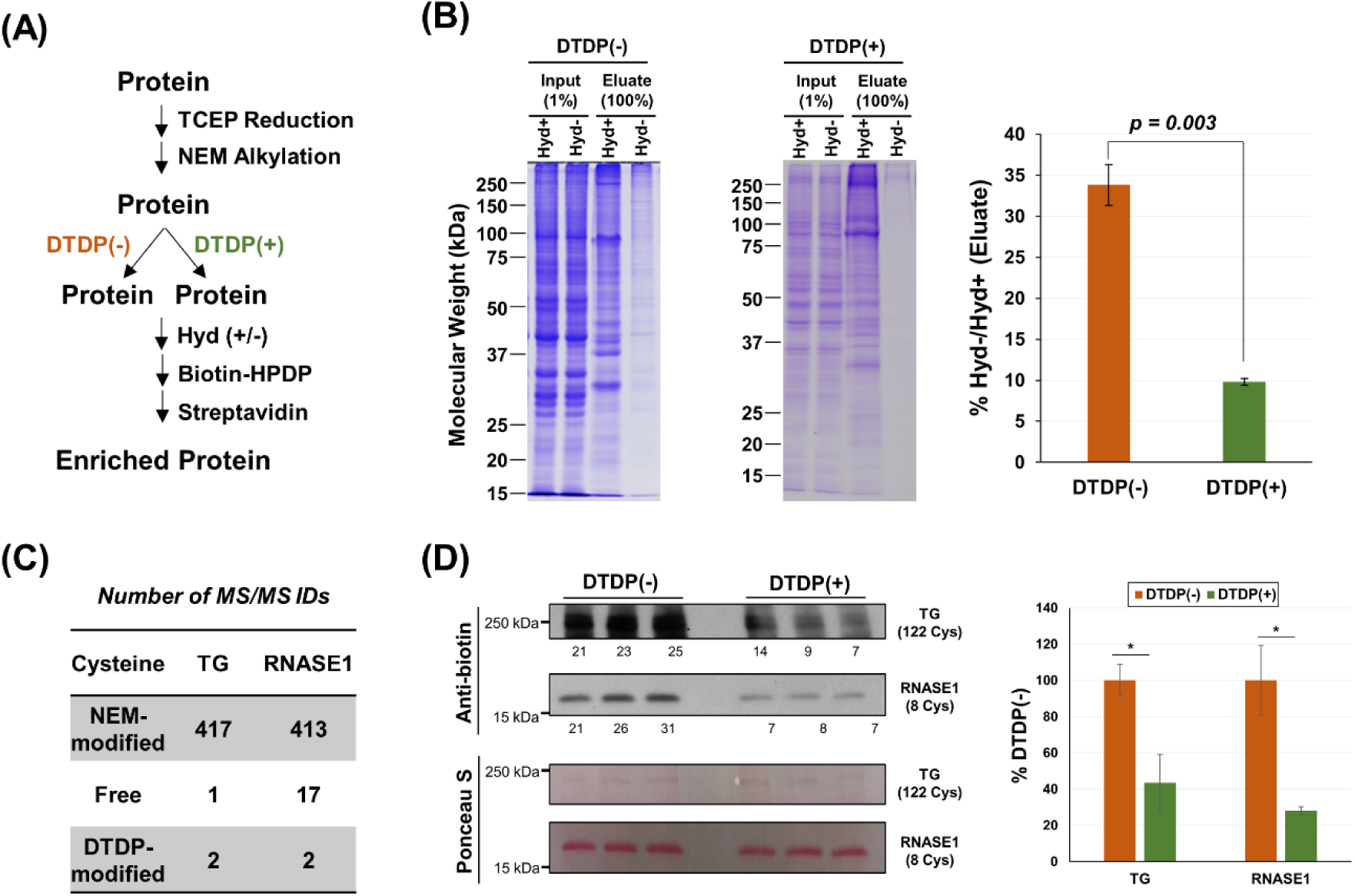
DTDP reaction largely eliminates the co-isolation of non-*S*-acylated proteins. **(A)** Workflow for the comparison of ABE enrichment with or without DTDP reaction. **(B)** Comparison of the levels of non-*S*-acylated proteins co-enriched by ABE, in the absence or presence of DTDP reaction. **(C)** Number of MS/MS identifications (*i.e.*, peptide-spectrum matches; PSMs) of cysteine-containing peptides derived from tryptic digestion of bovine thyroglobulin (TG) and ribonuclease A family member 1 (RNASE1) proteins. Protein mixture was reduced by TCEP, alkylated by NEM, further blocked by DTDP, digested with trypsin by FASP, and analyzed by 2D high-pH/low-pH LC-MS/MS. **(D)** Comparison of biotinylation levels of bovine TG and RNASE1 proteins with or without DTDP reaction. Protein mixture was reduced by TCEP, alkylated by NEM, treated with or without DTDP, reacted with biotin-HPDP, and subjected to immunoblotting analysis using anti-biotin conjugate of streptavidin-horseradish peroxidase. The numbers under blots indicate relative (% total) band intensities. * stands for *p* < 0.05.

To determine whether DTDP reacts with certain residual free cysteine residues (Fig. 3C), two commercially available and presumably non-*S*-acylated bovine proteins were used as model proteins: a) bovine thyroglobulin (TG), a 300-kDa protein containing 122 cysteine residues, was used to test whether DTDP reacts with many different residual cysteine residues or preferentially reacts with certain residual cysteine residues; and b) bovine ribonuclease A family member 1 (RNASE1), a 16-kDa protein containing 8 cysteine residues, was used to obtain high protein sequence coverage by LC-MS/MS, so the ratios of different modification forms (*i.e.*, NEM-modified, free, and DTDP-modified) can be determined for most if not all cysteine residues. The two proteins were mixed, reduced by TCEP, alkylated by NEM, reacted with DTDP, and digested with trypsin by FASP. Deep analysis by two-dimensional (2D) high-pH/low-pH LC-MS/MS suggested that a) NEM blocked the vast majority (95.6%-99.3%) of cysteine residues and b) DTDP selectively blocked certain residual free cysteine residues (Fig. 4C, Fig. S2, Table S4, and Table S5). Representative MS/MS spectra of DTDP-modified peptides were shown in Figures S3 and S4. In addition, as a positive control, bovine TG and RNASE1 protein mixture was reduced by TCEP, directly reacted with DTDP (*i.e.*, without NEM alkylation), digested by trypsin, and analyzed by LC-MS/MS. A total of 131 peptide-spectrum matches (PSMs) were identified from DTDP-modified TG or RNASE1 peptides with posterior error probability (PEP) values of <0.01 (Table S6), confirming that the DTDP modification was correctly configured for database searching.

To determine whether DTDP blockage prevents the biotin-HPDP reaction with non-*S*-acylated proteins, the TG and RNASE1 mixture was reduced by TCEP, alkylated by NEM, treated with or without DTDP, reacted with biotin-HPDP, and analyzed by immunoblotting to probe the biotinylation levels. As shown in Figure 4D, the DTDP blockage significantly reduced the biotinylation levels of both bovine proteins, compared with control samples without DTDP blockage. Collectively, further blockage of residual free cysteine residues by DTDP suppresses biotin-HPDP reaction, and thus largely eliminates the co-isolation of non-*S*-acylated proteins.

### LB-ABE Enables Deep *S*-acylproteomic Analysis

Compared with SILAC analysis, label-free proteomic analysis is less expensive and more scalable. Moreover, recent improvements of instrumentation and algorithms substantially increased the accuracy of label-free quantification (LFQ).^32^ Therefore, we coupled our LB-ABE method with label-free proteomics to profile the *S*-acylproteome of LNCaP cells (Fig. S5). To improve protein sequence coverage, three endoproteinases (*i.e.*, trypsin, Asp-N, and Arg-C) were used to digest LB-ABE-enriched proteins into peptides. For statistical comparison, three biological replicates were subjected to LB-ABE enrichment and endoprotease digestion, and each digestion product was analyzed by LC-MS/MS twice (Fig. S5).

A total of 4,025 proteins were identified with an FDR ≤ 1%, among which 3,531 protein groups were identified with at least two peptides (Table S7). For statistical comparison, among the 3,531 protein groups, 3,309 with at least three label-free quantification (LFQ) values in the Hyd(+) condition were selected (Table S8 and Fig. S6A). After imputation of the missing data and Student’s *t*-test followed by Benjamini-Hochberg correction, a total of 2,895 protein groups (87.5% of 3,309) were found to be significantly enriched (FDR<0.01, log_2_Ratio>1) in the Hyd(+) condition, compared with the control Hyd(-) condition (Table S8, Fig. S6B-C, and Fig. S7). To our knowledge, this represents so-far the largest set of high-confidence *S*-acylated protein candidates identified in a single study (n=2,895). It is about the same size of total *S*-acylated human proteins identified by different methods (n=2,881) and about 150% larger than the total size of human *S*-acylated proteins identified by ABE methods (n=1,173) (see Fig. 1A).

A comparison of the 2,895 candidate *S*-acylated proteins with the latest SwissPalm (v2) database (released on 02/18/2018) suggested that 1,571 (54.3% of 2,895) are known human *S*-acylated proteins (Table S8), confirming that our *S*-acylproteomic analysis enriched for *S*-acylated proteins. Based on the SwissPalm classification, of the 1,571 known *S*-acylated proteins, 123 are validated *S*-acylated proteins, 498 additional proteins are *S*-acylated protein candidates identified by two different *S*-acylproteomic techniques, and 950 additional proteins are *S*-acylated protein candidates identified by one *S*-acylproteomic technique (Fig. 5A). In addition to the 1,571 known *S*-acylated proteins, 1,324 novel *S*-acylated protein candidates were discovered by this study (Fig. 5A).

**Figure 5.**
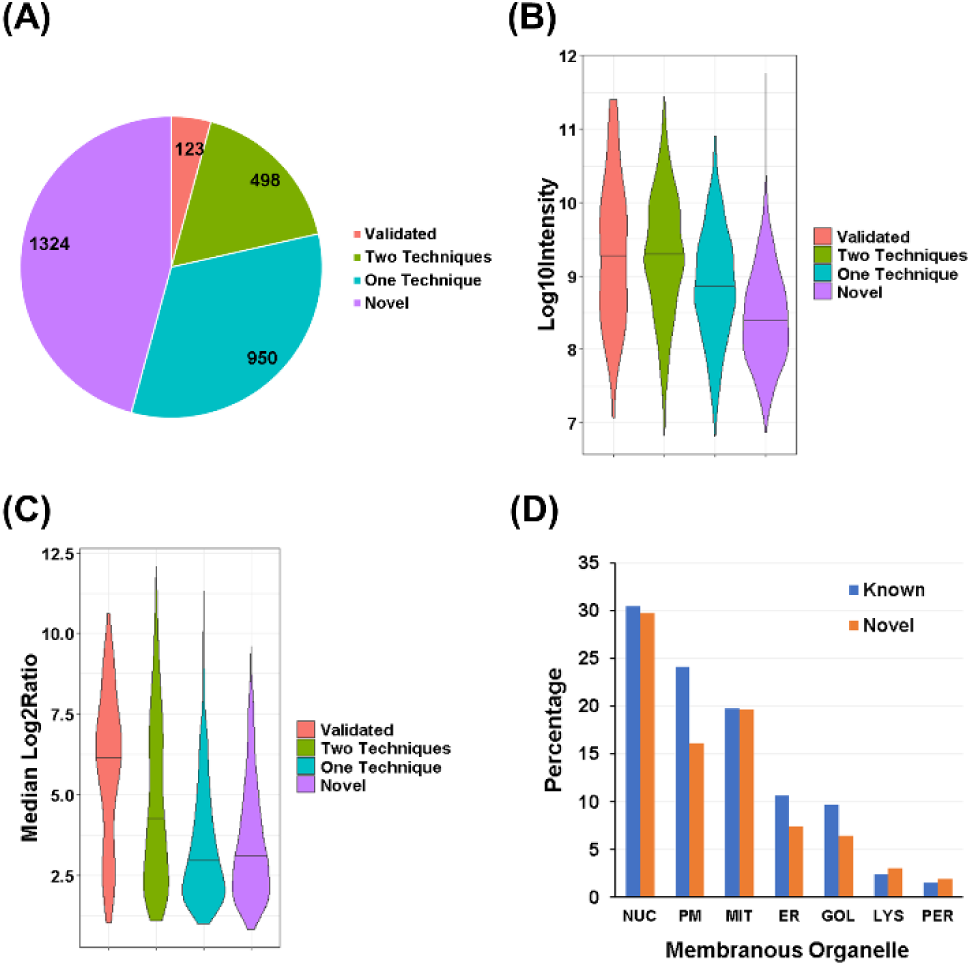
Comparison of the known and novel *S*-acylated proteins. **(A)** Pie chart of the 2,895 candidate *S*-acylated proteins, according to the SwissPalm classification. **(B)** Violin plot of the 2,895 candidate *S*-acylated proteins according to the log_10_-transformed total ion intensities. **(C)** Violin plot of the 2,895 candidate *S*-acylated proteins according to the median log_2_-transformed Hyd+/Hyd− ratios. **(D)** Histogram of known and novel candidate *S*-acylated proteins based on their subcellular localizations.

As mentioned above, the LB-ABE method largely eliminates the co-isolation of non-*S*-acylated proteins. Consistently, a comparison of the log_2_-transformed Hyd+/Hyd− ratios suggested that the LB-ABE method provided much cleaner enrichment (*i.e.*, higher Hyd+/Hyd− ratios) than the ABE method (Fig. S8). Moreover, the significant reduction of non-*S*acylated protein co-enrichment may alleviate the masking effect and thus substantially improve the detection of lowly abundant *S*-acylated proteins―the low abundance may be due to low protein expression levels, low *S*-acylation levels, or both. Consistently, compared with the three groups of known human *S*-acylated proteins, the novel candidate *S*-acylated proteins are generally less abundant and have significantly lower log10-transformed total ion intensities (Fig. 5B and Fig. S9A). Nonetheless, the fold-enrichment of the novel candidates are at least comparable to the known *S*-acylated proteins identified by one technique (Fig. 5C and Fig. S9B). In addition, according to the gene ontology (GO) annotation, the subcellular distribution of the novel *S*-acylated protein candidates is similar to that of the known *S*-acylated proteins (Fig. 5D). The major difference is in the plasma membrane (PM), the endoplasmic reticulum (ER), and the Golgi apparatus (GOL), where the numbers of novel *S*-acylated protein candidates are about 30% less than those of known *S*-acylated proteins (Fig. 5D). This is likely because almost all human PATs are localized in the three organelles,^33^ the *S*-acylation levels of proteins resident in these three organelles might be relatively high, so they tend to be more easily identified in previous studies. In addition, since the release of the SwissPalm (v2) database on 02/18/2018, some new *S*-acylation studies, including global *S*acylproteomic studies, were published and added to the SwissPalm database. Mapping of the 1,324 novel *S*-acylated protein candidates against the online SwissPalm database (as of 03/18/2019) suggested that 20 proteins were identified as (candidate) *S*-acylated proteins by independent studies in the past year (Table S7), further corroborating our *S*-acylproteomic findings.

### Validation of Novel Candidate *S*-acylated Proteins

Interestingly, a number of proteins important for PCa development and progression were identified as high-confidence candidate *S*-acylated proteins in LNCaP cells. These include known *S*-acylated proteins such as AR, CTNNB1, EGFR, ERBB2, and EZH2 as well as novel *S*-acylated protein candidates such as AMACR, FOLH1 (also known as prostate-specific membrane antigen, PSMA), FOXA1, KLK3 (also known as prostate-specific antigen, PSA), and TMPRSS2.

To validate the findings, *S*-acylated proteins were enriched from three biological replicates of LNCaP cells, followed by immunoblotting analysis of known or candidate *S*-acylated proteins. As shown in Fig. 6A and Fig. 6B, after LB-ABE enrichment, all the tested proteins were highly enriched in the Hyd+ condition, compared with the Hyd− condition, confirming the *S*-acylproteomic findings. To exclude the possibility that the apparent enrichment might arise from different recovery efficiency of proteins under different (*i.e.*, Hyd+ vs Hyd−) conditions, an anti-actin primary antibody (a presumably non-*S*-acylated protein) was spiked into LNCaP cell lysates. Fig. 6C indicates that, after LB-ABE enrichment, similar amounts of residual anti-actin antibody protein were detected in both Hyd+ and Hyd− conditions, confirming that protein recovery efficiencies for both conditions were similar. In addition, densitometric analysis of immunoblots from eluate (*i.e.*, enriched) versus input samples suggested that the estimated *S*-acylation levels of these PCa-related proteins were generally low, ranging from 0.6% to 1.8% (Fig. 6A-B and Fig. S10). To exclude the possibility that the detected low *S*-acylation levels were due to low recovery of *S*-acylated proteins, a well characterized *S*-acylated protein caveolin-1 (CAV1) was used as a positive control. Because CAV1 is not expressed in LNCaP cells but is expressed in PC3 cells, the *S*-acylation level of CAV1 was estimated in PC3 cells. As shown in Fig. 6D, the *S*-acylation level of CAV1 was about 100%, confirming that our LB-ABE method provides high recovery of *S*-acylated proteins.

**Figure 6.**
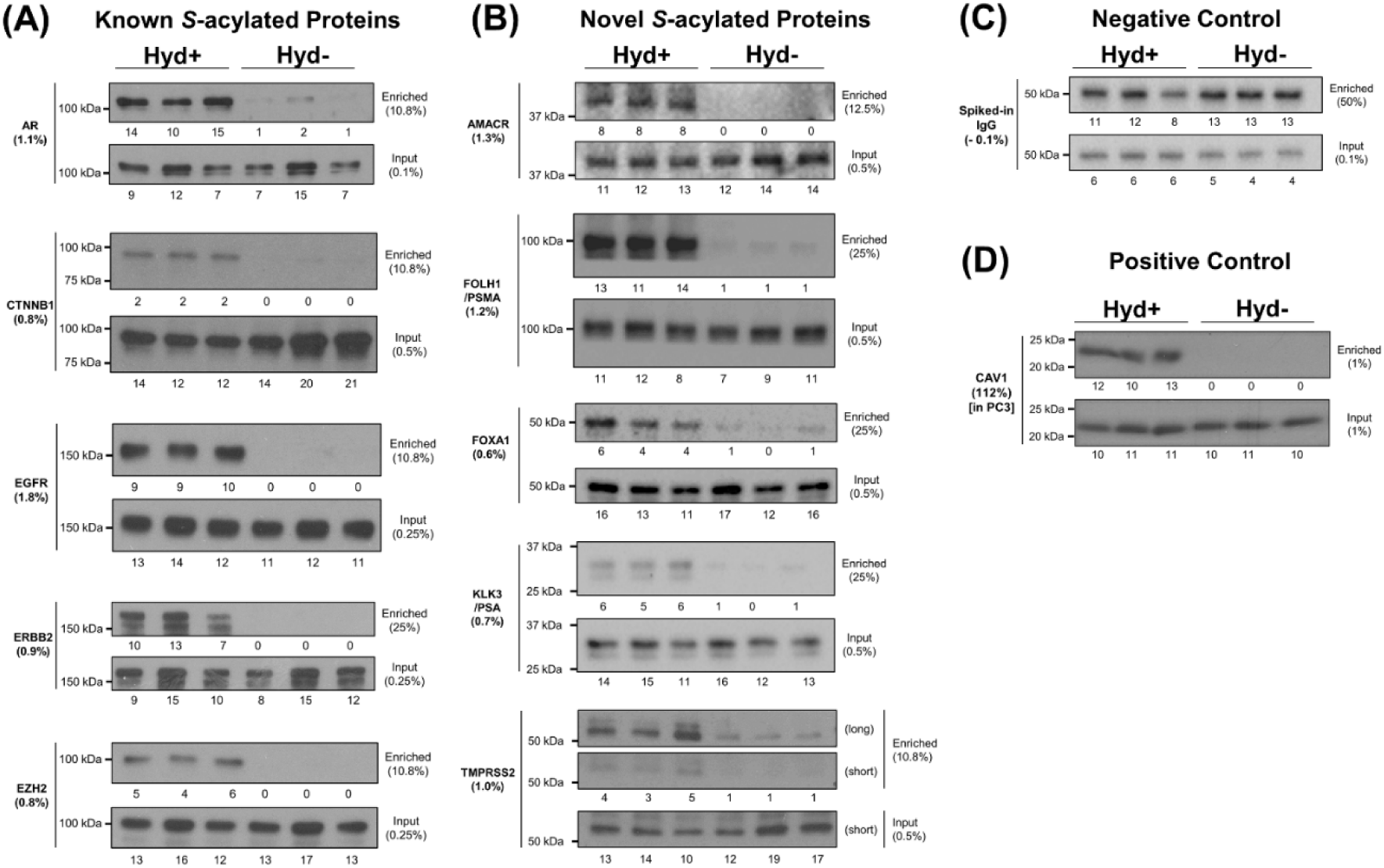
Validation of candidate *S*-acylated proteins and estimation of *S*-acylation levels by immunoblotting. **(A)** Immunoblotting analysis of known *S*-acylated proteins in LNCaP cells. **(B)** Immunoblotting analysis of novel *S*-acylated protein candidates in LNCaP cells. **(C)** Immunoblotting analysis of a non-*S*-acylated protein (anti-actin antibody) spiked into LNCaP cell lysates as a control to ensure that there is no significant difference of protein recovery under the Hyd+ and Hyd− conditions. **(D)** Immunoblotting analysis of caveolin-1 (CAV1) in PC3 cells as a positive control demonstrating the high recovery of *S*-acylated proteins offered by the LB-ABE method. In the figure, the input samples were taken from protein solution prior to streptavidin affinity purification, and the enriched samples were proteins eluted from streptavidin agarose beads by TCEP. The numbers below gene names indicate *S*-acylation levels estimated from the comparison of band densities from “Enriched” and “Input” samples. The numbers below blots indicate relative (% total) band densities. The numbers below “Enriched” and “Input” show the percentages of samples used for the immunoblotting analysis. See Fig. S10 for uncropped blots.

### Limitations of the Study

Like all other ABE methods, the LB-ABE method requires multiple methanol/chloroform precipitation steps, which presumably may cause gradual sample loss. However, we found that methanol/chloroform precipitation provided >97% protein recovery (Fig. S11) and that the total protein loss after eight rounds of methanol/chloroform precipitations was <20%. Currently, we are developing a simpler and more streamlined method to eliminate the time-consuming and labor-intensive protein precipitation and re-solubilization steps.

Another caveat of the study is that no *S*-acylation sites were mapped. However, because *S*-acyl groups are removed by Hyd during ABE or LB-ABE enrichment, LC-MS/MS analysis only provide putative *S*-acylation sites. Currently, we are developing a new technology for proteome-scale identification of intact *S*-acylated peptides, which will provide definitive information of not only *S*-acylation sites but also the fatty *S*-acyl species covalently attached to the sites.

## CONCLUSIONS

In summary, the LB-ABE method largely eliminates the co-isolation of non-*S*-acylated proteins, provides much higher signal-to-noise ratios, and enables deep *S*-acylproteomic profiling. The comprehensive *S*-acylproteomic analysis of LNCaP cells identified nearly 3,000 high-confidence candidate *S*-acylated proteins, including about 1,600 known *S*-acylated proteins. Immunoblotting analysis of LB-ABE enriched proteins suggested that many PCa-related proteins are *S*-acylated at a low (∼1%) level in regularly cultured LNCaP cells. The LB-ABE method is expected to facilitate more comprehensive identification and more accurate quantification of *S*-acylated proteins across different conditions in various biological samples, such as tissue specimens and extracellular vesicles. In addition, the LB-ABE can potentially be adapted for low-background enrichment analysis of many other cysteine modifications like *S*-nitrosylation.

## ASSOCIATED CONTENT

### Supporting Information

Figures S1-11 (PDF)

## AUTHOR INFORMATION

### Author Contributions

WY conceived and led the project. BZ, YW, YY and JM performed the experiments and analyzed the data. BZ and WY wrote the manuscript. DDV, MRF and WY discussed the results and edited the manuscript. All authors have given approval to the final version of the manuscript.

### Notes

The authors declare no competing financial interest.

## ACKNOWLEDGMENT

We gratefully acknowledge financial support from the Department of Defense (W81XWH-15-1-0167 to WY), the Spielberg Family Team Science Fund (CSR205927 to MRF), and the National Cancer Institute (5R01CA218526 to DDV and WY).

