## Supplemental Figures for "Low-background Acyl-biotinyl Exchange Largely Eliminates the Co-isolation of Non-S-acylated Proteins and Enables Deep S-acylproteomic Analysis"

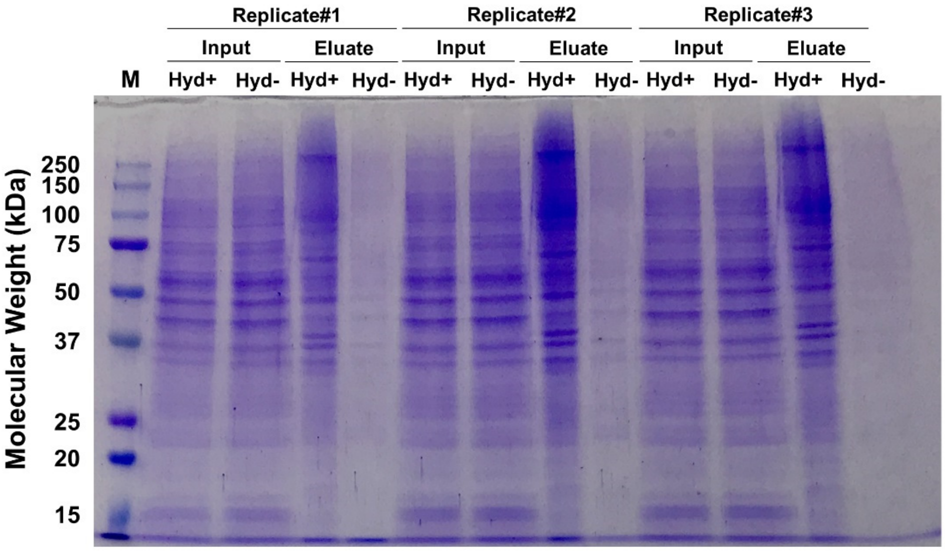


**Figure S1.** SDS-PAGE image of input and enriched proteins from human prostate cancer PC3 cells. Three biological replicates of PC3 cells were subjected to LB-ABE enrichment. "Input" and "Eluate" samples indicate proteins before and after streptavidin affinity purification, respectively. Hyd: Hydroxylamine.

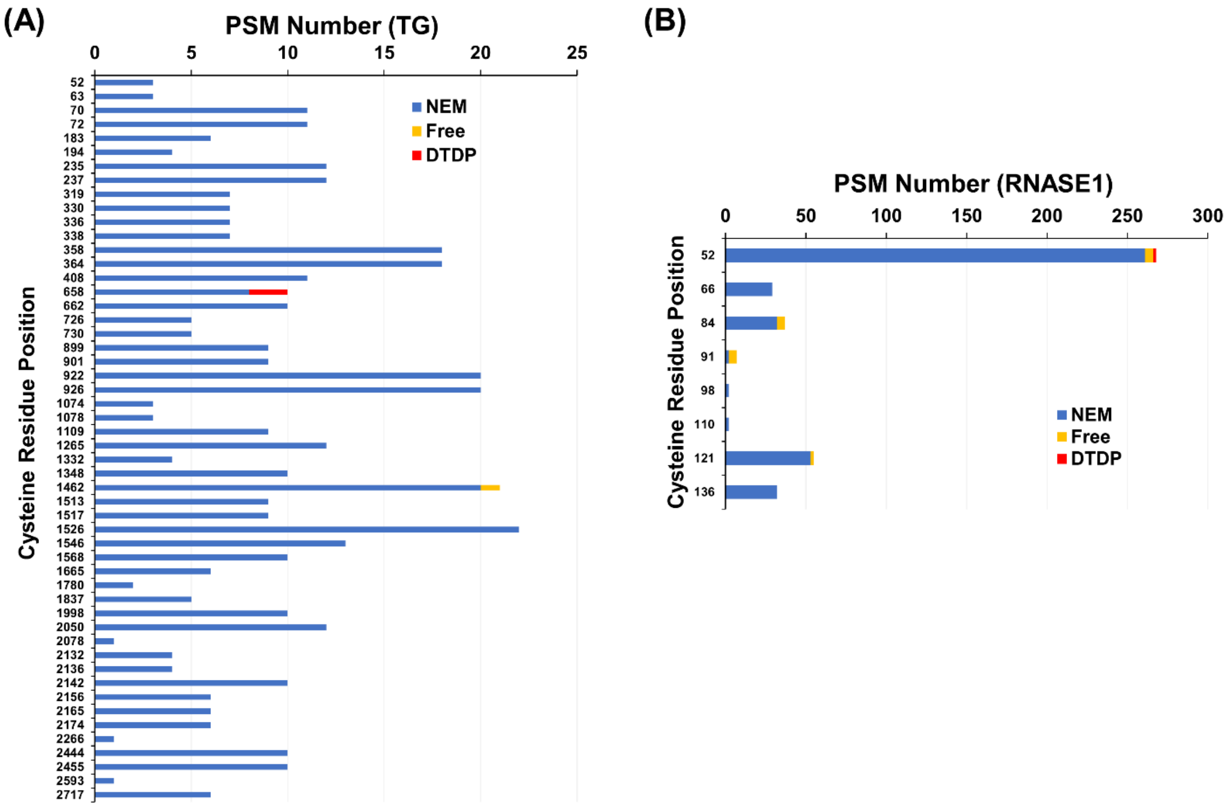


**Figure S2.** Bar graphs of peptide-spectrum-match (PSM) numbers identified for each cysteine residue of bovine **(A)** thyroglobulin (TG) and **(B)** ribonuclease A family member 1 (RNASE1). Only PSMs identified with posterior error probabilities (PEPs) of <0.01 were used. NEM: N-ethylmaleimide; DTDP: 2,2'-dithiodipyridine.

| Raw File | Scan | Method | Score | m/z |
| --- | --- | --- | --- | --- |
| Bovine_bRPLC_Fr10 | 9795 | FTMS; HCD | 107.67 | 522.69 |

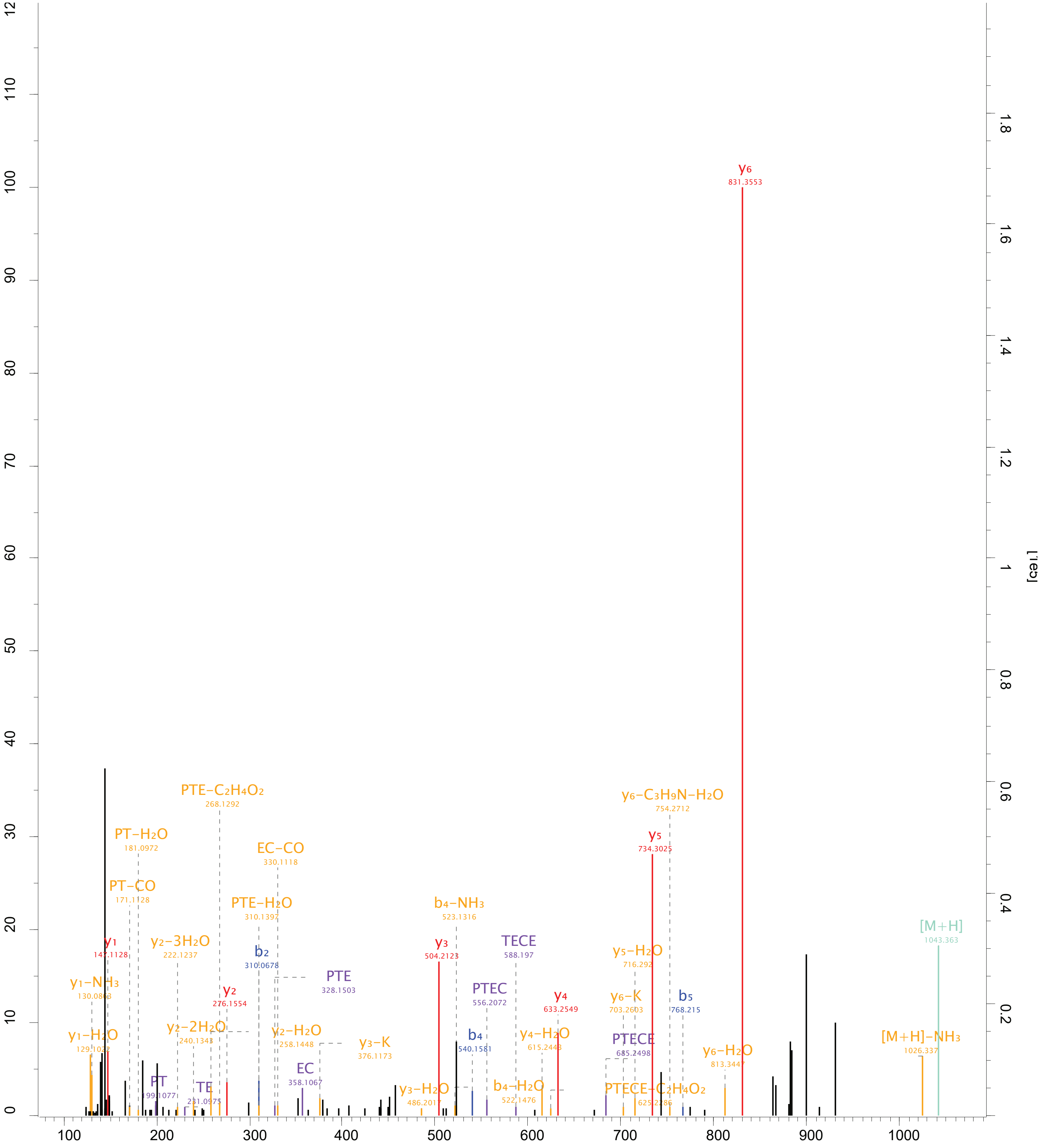


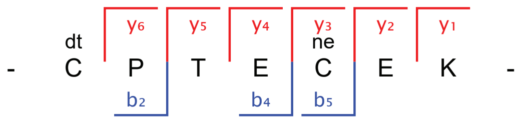

**Figure S3.** A representative MS/MS spectrum of DTDP-modified peptide from bovine thyroglobulin (TG). The spectrum was generated using the Expert Annotation algorithm in the MaxQuant (v1.5.5.1) environment.

| Raw File | Scan | Method | Score | m/z |
| --- | --- | --- | --- | --- |
| Bovine_bRPLC_Fr13 | 9733 | FTMS; HCD | 95.16 | 816.97 |

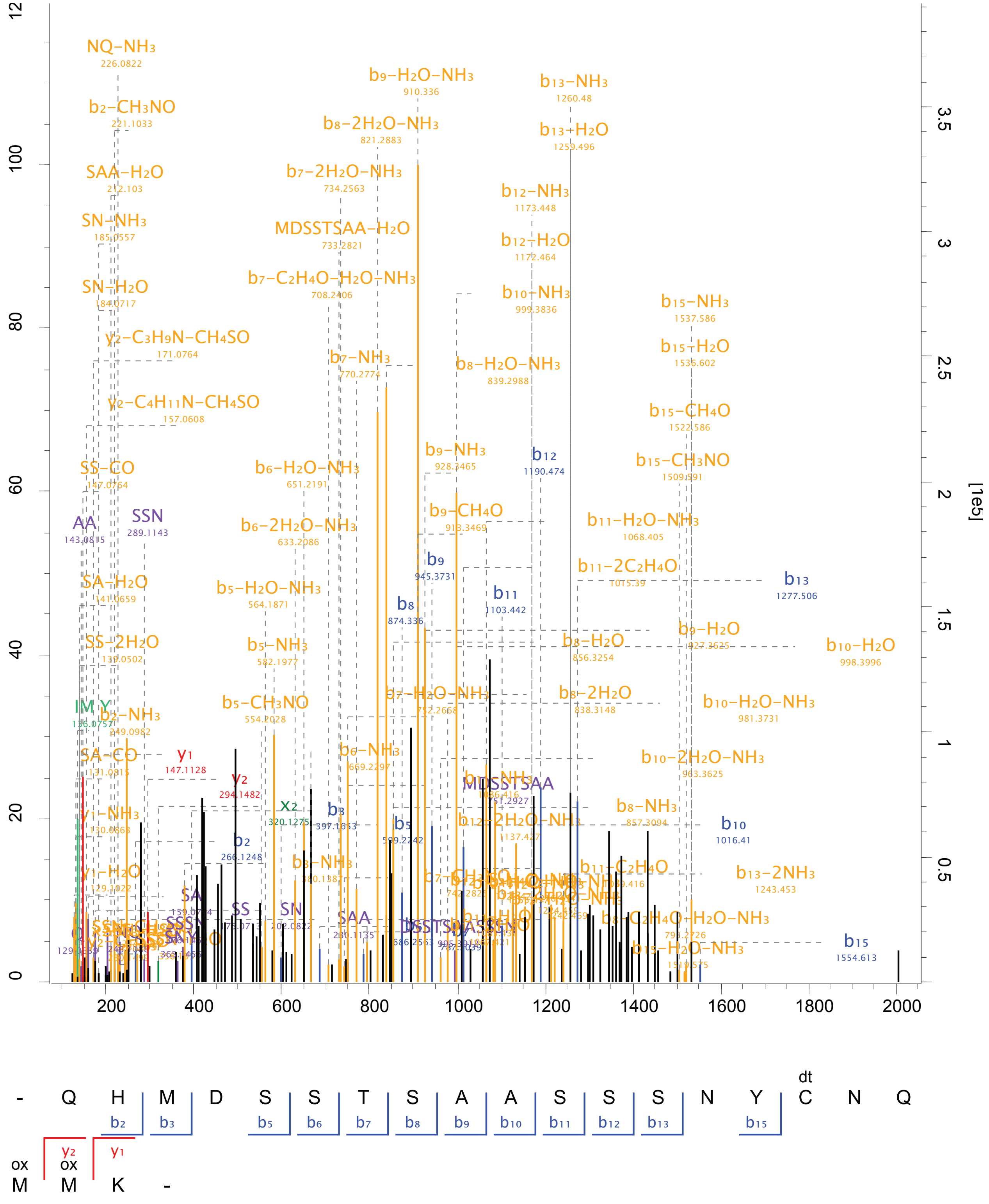

**Figure S4.** A representative MS/MS spectrum of DTDP-modified peptide from bovine ribonuclease A family member 1 (RNASE1). The spectrum was generated using the Expert Annotation algorithm in the MaxQuant (v1.5.5.1) environment.

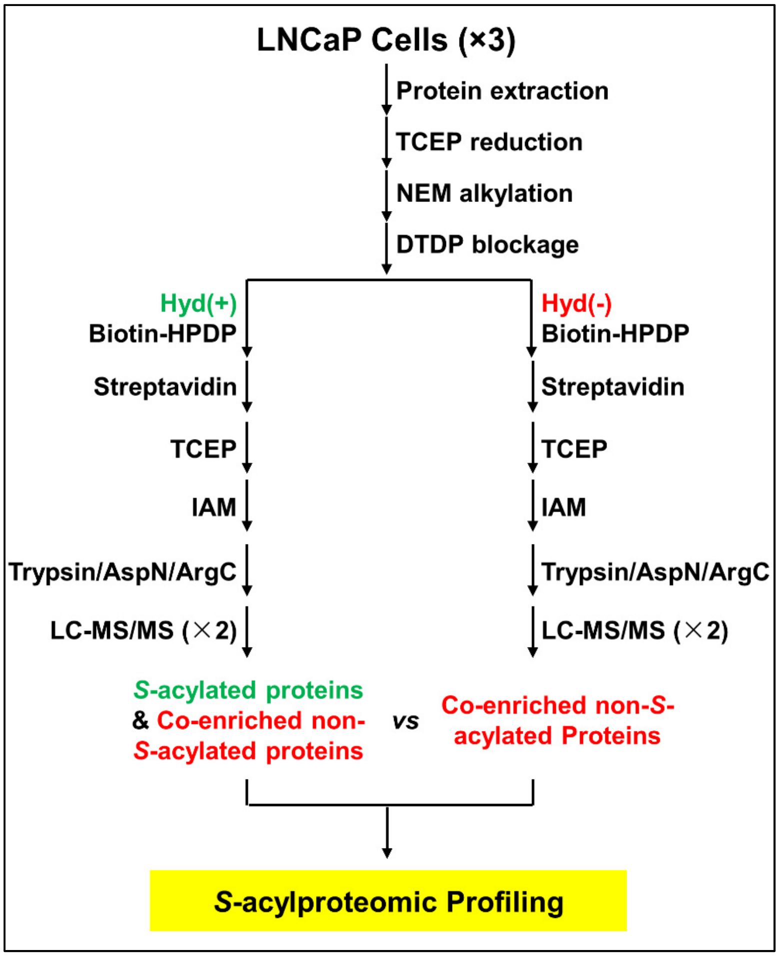


**Figure S5.** Workflow for the *S*-acylproteomic profiling of human LNCaP cells by coupling LB-ABE with label-free quantitative proteomics. TCEP: tris(2-carboxyethyl)phosphine; NEM: N-ethylmaleimide; DTDP: 2,2'-dithiodipyridine; Hyd: hydroxylamine; IAM: iodoacetamide.

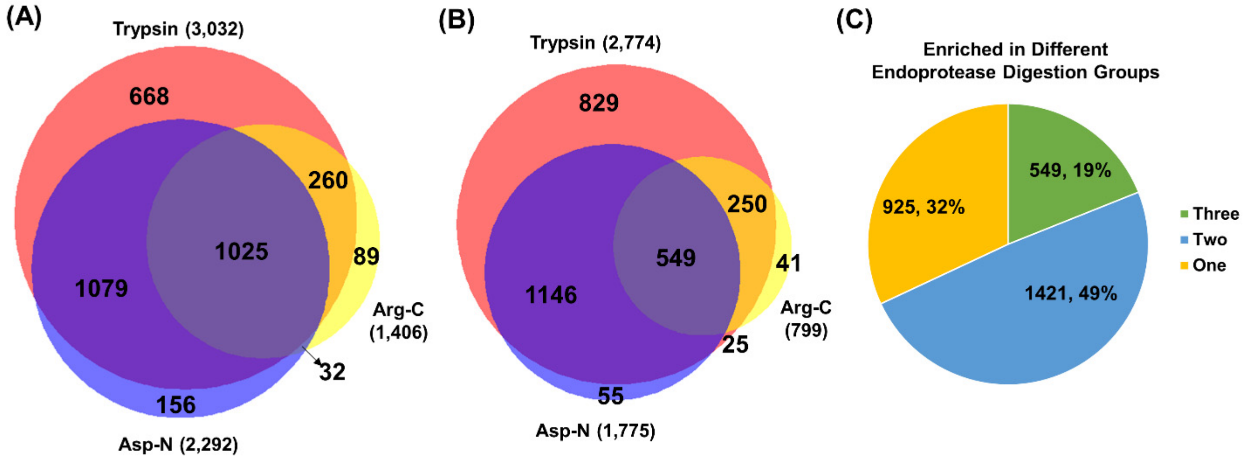


**Figure S6.** Comparison of protein numbers obtained by digestion with three different endoproteases (*i.e.*, trypsin, Asp-N, and Arg-C). **(A)** Venn diagram of proteins (out of 3,309 in total) identified from each endoprotease digestion group. **(B)** Venn diagram of significantly ($q$<0.01, $\log_2$ratio>1) enriched *S*-acylprotein candidates (out of 2,895 in total) identified from each endoprotease digestion group. **(C)** Pie chart of the 2,895 significantly enriched *S*-acylprotein candidates, which were classified based on the number of endoprotease digestion groups (*i.e.*, shared by three enzyme groups, shared by two enzyme groups, and only by one enzyme).

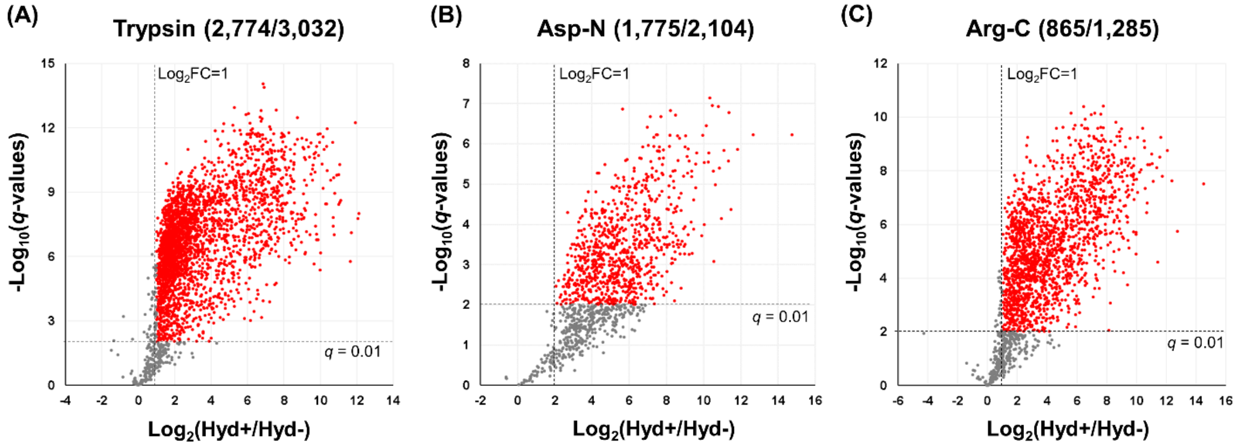


**Figure S7.** Volcano plots of significance ($\log_{10}$-transformed $q$ values) versus fold change ($\log_2$-transformed ratios) for proteins quantified from **(A)** trypsin, **(B)** Asp-N, and **(C)** Arg-C digestion groups. Red dots show significantly ($q$<0.01 and $\log_2$Ratio>1) enriched proteins. Each dot represents a protein.

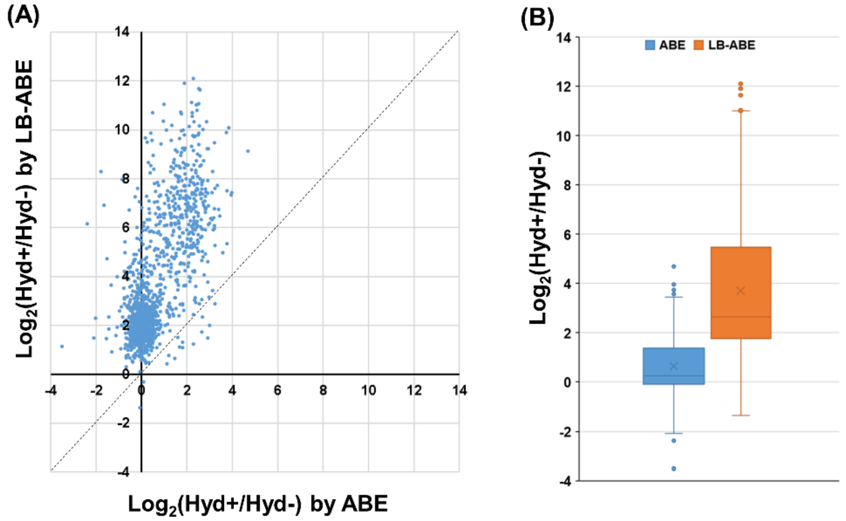


**Figure S8.** Comparison of $log_2$-transformed Hyd+/Hyd- ratios obtained using ABE versus LB-ABE for the overlapping set (n=1,315) of proteins identified from trypsin digestion groups. **(A)** Scatter plot of $log_2$ratios. The diagonal dotted line represents a 1:1 ratio. Each dot represents a protein. **(B)** Box-plot comparison of $log_2$ratios obtained using ABE versus LB-ABE for the overlapping set (n=1,315) of proteins.

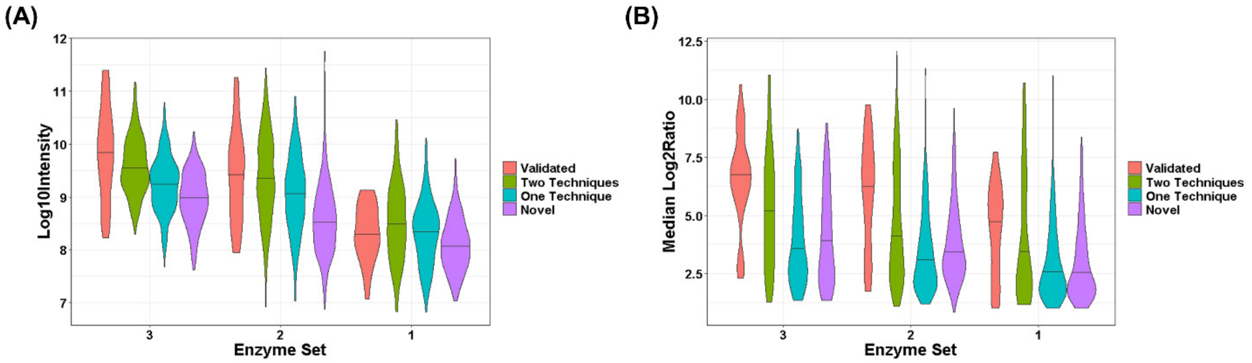


**Figure S9.** Violin plots of significantly enriched proteins identified from different number of endoprotease digestion groups (*i.e.*, shared by three enzyme groups, shared by two enzyme groups, and only by one enzyme). **(A)** Violin plot of significantly enriched proteins according to their $log_{10}$-transformed total ion intensities. **(B)** Violin plot of significantly enriched proteins according to their $log_2$-transformed Hyd+/Hyd- ratios (median values of three digestion groups). Significantly enriched proteins were classified based on SwissPalm (v2) annotations: 1) validated *S*-acylated proteins, 2) candidate *S*-acylated proteins identified by two independent techniques, 3) candidate *S*-acylated proteins identified by one technique, and 4) novel candidate *S*-acylated proteins.

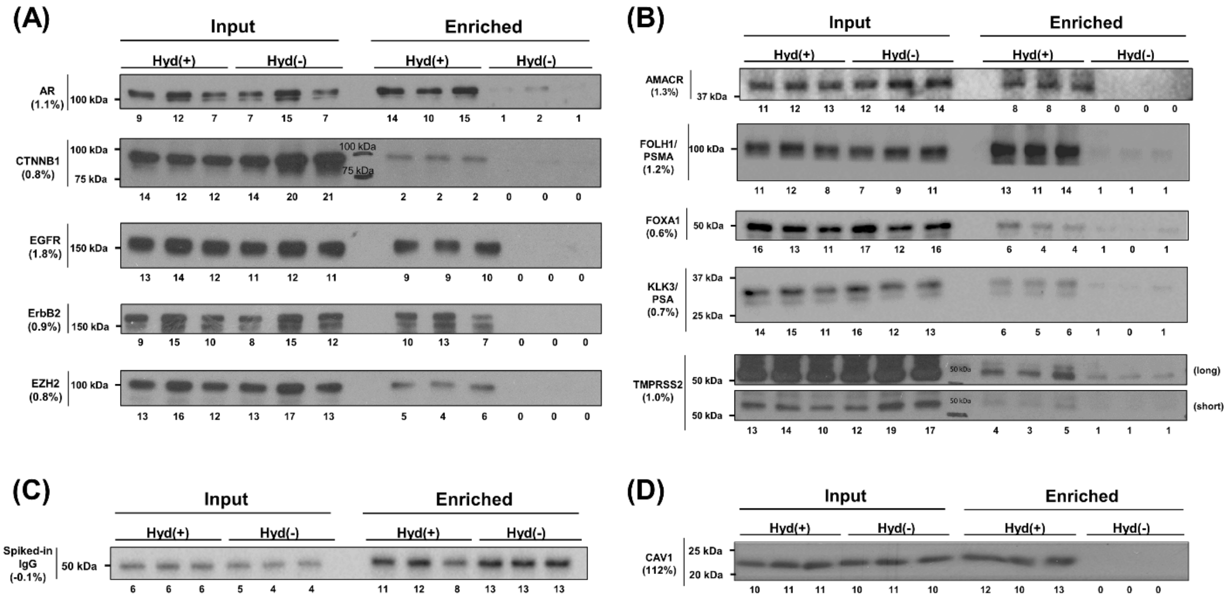


**Figure S10.** Validation of candidate *S*-acylated proteins and estimation of their *S*-acylation levels. **(A)** Immunoblotting analysis of known *S*-acylated proteins in LNCaP cells. The percentages of samples used for the blots were (in the order of "Enriched" and "Input"): AR (10.8%, 0.1%), CTNNB1 (10.8%, 0.5%), EGFR (10.8%, 0.25%), ERBB2 (25%, 0.25%), and EZH2 (10.8%, 0.25%). **(B)** Immunoblotting analysis of novel *S*-acylated protein candidates in LNCaP cells. The percentages of samples used for the blots were (in the order of "Enriched" and "Input"): AMACR (12.5%, 0.5%), FOLH1/PSMA (25%, 0.5%), FOXA1 (25%, 0.5%), KLK3 (25%, 0.5%), and TMPRSS2 (10.8%, 0.5%). **(C)** Immunoblotting analysis of a presumably non-*S*-acylated protein (anti-β-actin antibody) spiked into LNCaP cell lysates as a control, in order to ensure that there was no significant difference of protein recovery under the Hyd+ and Hyd- conditions. 50% of "Enriched" sample and 0.1% of "Input" sample were used for the immunoblotting experiment. **(D)** Immunoblotting analysis of caveolin-1 (CAV1) in PC3 cells as a positive control demonstrating the high recovery of *S*-acylated proteins offered by the LB-ABE method. 1% of "Enriched" and 1% of "Input" sample was used for the immunoblotting experiment. In the figure, the input samples were taken from protein solution prior to streptavidin affinity purification, and the enriched samples were proteins eluted from streptavidin agarose beads by TCEP. The numbers below gene names indicate *S*-acylation levels estimated from the comparison of band densities from "Enriched" and "Input" samples. The numbers below blots indicate relative (% total) band densities.

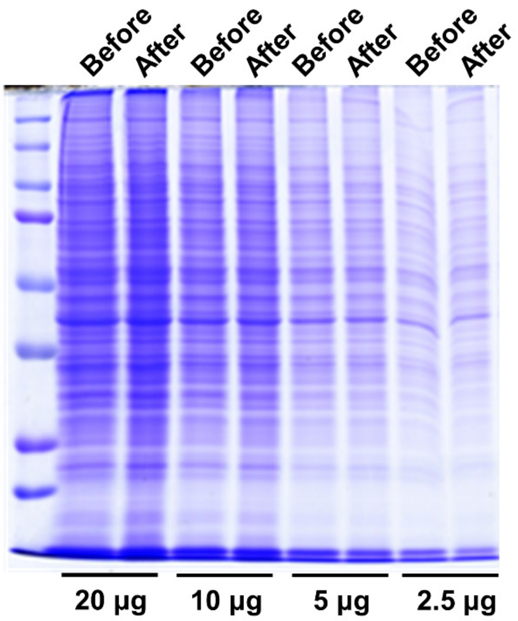


**Figure S11.** SDS-PAGE analysis of protein recovery efficiency of methanol/chloroform precipitation. The protein recovery was nearly 100% for different amounts (2.5-20 μg) of input proteins after methanol/chloroform precipitation.
